# Intranasal delivery of a broadly neutralizing single domain antibody targeting ACE2 protects against SARS-CoV-2 infection

**DOI:** 10.1101/2024.10.11.617877

**Authors:** Simon Blachier, Ignacio Fernández, Laurine Conquet, Emilie Giraud, Isabelle Staropoli, Vincent Michel, Fruzsina Szilagyi, Salomé Guez, Alix Boucharlat, Jeanne Chiaravalli, Jaouen Tran-Rajau, Evelyne Dufour, Stéphane Petres, Delphine Planas, Xavier Montagutelli, Fabrice Agou, Felix Rey, Pierre Lafaye, Jost Enninga, Gabriel Ayme, Olivier Schwartz, Anne Brelot

## Abstract

SARS-CoV-2 accumulates mutations over time leading to the emergence of variants, which become largely resistant to existing vaccines and spike protein-targeted antiviral treatment. Therefore, there is a need for other therapies with broad efficiency. Here, we targeted the angiotensin-converting enzyme 2 (ACE2), the major entry receptor for SARS-CoV-2. We purified three single domain heavy chain antibodies (VHHs) after immunization of an alpaca with the ectodomain of ACE2. These VHHs bound ACE2 with nanomolar affinity and specifically detected membrane-anchored ACE2. Two of them (B07 and B09) neutralized by a competitive mechanism multiple SARS-CoV-2 isolates, including Omicron variants (XBB.1.16.1; EG.5.1.3; BA.2.86.1), without impacting the proteolytic activity of the enzyme. Fusion of B07 with conventional Fc domain markedly improved its binding and neutralizing efficacy. This dimeric Fc-conjugated B07 (B07-Fc) recognized specific residues of the N-terminal helix 1 of ACE2. When administrated prophylactically and intranasally, B07-Fc induced a strong dose-dependent protection of mice expressing human ACE2 (K18-hACE2) from SARS-CoV-2 Omicron. Hamsters were weakly protected due to low binding of B07-Fc to hamster ACE2. These single domain antibodies targeting hACE2 represent potential broad-spectrum therapeutic candidates against any emerging viruses using ACE2 as a receptor. These inhalable neutralizing single domain antibodies also represent a non-invasive approach against respiratory viral infection.

## Introduction

Severe Acute Respiratory Syndrome Coronavirus 2 (SARS-CoV-2) is the causative agent of COVID-19 (coronavirus disease 2019). So far, it has infected more than 760 million people with more than 6.9 million deaths worldwide, even though these numbers are likely underestimated (World Health Organization - WHO, 2023). To date, multiple COVID-19 vaccines, effective against severe forms of COVID-19, have been authorized by the WHO, making it possible to control the pandemic. However, the emergence of new variants represents a continuing threat of resurgence^1^. Indeed, depending on the SARS-CoV-2 variants, the vaccines differ in their protection against transmission, and they lose their effectiveness over time^2–4^. Hence, constant, and rapid updating of vaccines is critical but remains a challenge. Vaccine protection may also be insufficient in individuals with altered immune status^5^. For these reasons, passive immunization with monoclonal antibodies neutralizing SARS-CoV-2 spike (S) protein has been developed and used as therapeutic and prophylactic intervention against infection^6–8^. However, their effectiveness differs according to the variants, and they poorly protect against the latest Omicron variants^1,4–6,9^. Therefore, other strategies are needed to complete the arsenal against COVID-19.

SARS-CoV-2 infects cells by interacting with a receptor: the angiotensin-converting enzyme 2 (ACE2), and then by driving fusion of the viral membrane and the cell membrane. Both processes, receptor binding and membrane fusion, are mediated by the S protein present at the surface of the virus^10^. ACE2 is a type I integral membrane glycoprotein widely expressed in almost all tissues^11^, which regulates the Renin-Angiotensin-System^12^. Besides its cellular form, ACE2 exists as a cleaved and enzymatically active form in the plasma^13^. Soluble ACE2 may indirectly be involved in SARS-CoV-2 entry ^14^. Using soluble derivatives of ACE2 as a “decoy” to trap the S protein is one strategy being studied to avoid viral escape^15–17^.

Here, we developed a strategy for the direct targeting of trans-membrane ACE2 instead of S. Because of the major role of ACE2 in regulation of blood pressure, and in vascular, renal, and myocardial physiology, the challenge was to develop compounds, which target S binding surface on ACE2 without inhibiting the catalytic activity of this enzyme. The enzyme’s active site and S-binding surface are distinct domains of ACE2, suggesting that this is feasible^18,19^. Monoclonal antibodies targeting ACE2 proved their efficacy in inhibiting SARS-CoV-2 infection of multiple variants^20–23^. An alternative to mAbs for clinical application is the use of camelid single domain antibodies (also called single domain VHH antibody or VHH), which correspond to the variable region of camelid heavy-chain antibody. These fragments, 10 times smaller than « conventional » antibodies, retain full antigen-binding capacity, have a propensity to bind small epitopes, and are easily produced in prokaryotic expression systems^24^. Moreover, VHHs are amenable to covalent linkage with each other or with the Fc domain for example, thereby increasing their valency and stability, and improving their pharmacodynamic profil *in vivo*^25–27^. Therefore, they can be delivered to the lungs by inhalation, which can confer significant advantage for SARS-CoV-2 treatment. These types of approach have been used successfully to treat other diseases. For example, a VHH-based therapy was approved in the USA and in Europe in 2019 for the treatment of acquired thrombotic trombocytopenic purpura^28^.

In this study, we produced monomeric and dimeric Fc-conjugated VHHs that (1) specifically bind ACE2 with nanomolar affinity, (2) recognize ACE2 at the surface of different cell types, (3) do not alter the proteolytic activity of ACE2 both *in vitro* and *in cellulo*, and (4) efficiently neutralize multiple variants of SARS-CoV-2 *in vitro* (Delta, BA.1, BQ.1.1, XBB.1.5, XBB.1.16.1, EG.5.1.3; BA.2.86.1). These VHHs compete S binding by targeting residues in the N-terminal helix 1 of ACE2. Moreover, Fc-conjugated VHHs are active in mouse K18-hACE2 and hamster against Omicron after intranasal inoculation. Therefore, these VHHs are promising candidates with broad neutralizing potential against any emerging variants of SARS-CoV-2 following respiratory administration.

## Results

### Production of VHHs targeting human ACE2

We immunized an alpaca with the soluble catalytic ectodomain of human ACE2 (sACE2). From lymphocytes, we constructed a cDNA library containing about 8×10^7^ different VHHs sequences. We selected the produced VHHs by phage display through three rounds of panning against sACE2, each with different buffer and washing conditions. In total, 192 individual clones were assayed by ELISA to test whether they could bind sACE2. Three specific VHHs were obtained, called B07, B09, and B10. B10 presented a substantially different sequence in their variable domains (CDRs) compared to B07 and B09. B07 and B09 differed by only 6 amino-acid mutations in their CDRs.

### VHHs B07, B09, and B10 interact with sACE2 in a nanomolar range

The different VHHs were expressed in fusion with His and c-myc tags at the C terminal end. We characterized the interaction between His-VHH and sACE2 by determining association (k_on_) and dissociation (k_off_) rate constants using biolayer interferometry (BLI) technology (Octet HTX, Sartorius)^29^ (**Fig. 1A**). We immobilized His-VHH on Ni-NTA biosensors and incubated them with different concentrations of sACE2. From these data, we deduced a K_D_ value of 350 nM, 353 nM and 10 nM for B07, B09, and B10, respectively (**Fig. 1A**), indicating that these VHHs bind sACE2 with good (B07, B09) and high (B10) affinity.

**Fig. 1:**
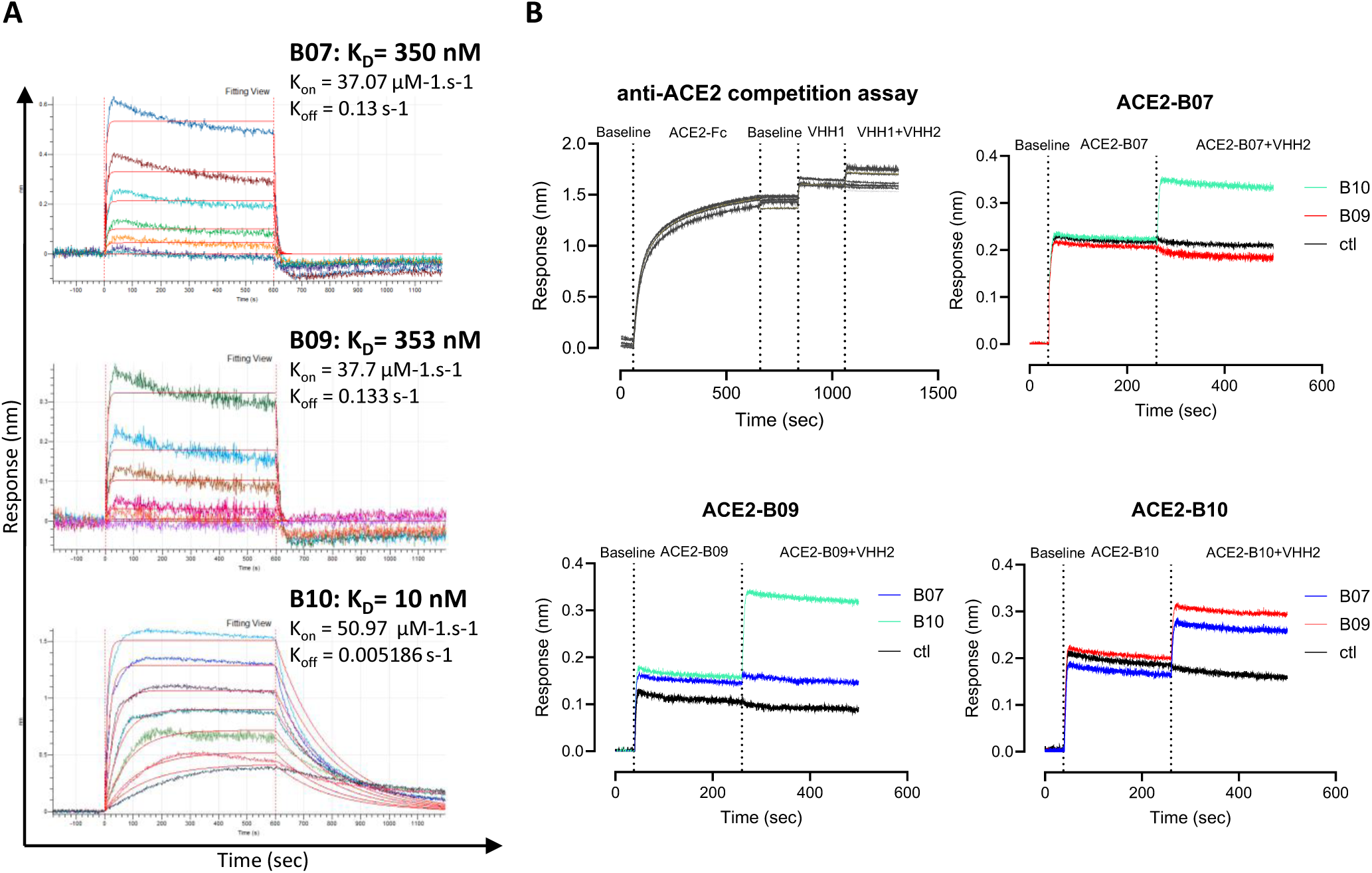
VHHs binding on sACE2. **(A)** Kinetic analysis of sACE2 on His-VHH-B07, His-VHH-B09, and His-VHH-B10 by BioLayer Interferometry (BLI) using various concentrations of sACE2 (0.3 ; 0.625 ; 1.25 ; 2.5 ; 5 ; 10 ; 20 µg/ml). Ni-NTA biosensors were used to immobilize 5 µg/ml of His-VHH. K_D_ of each VHHs are indicated. One representative experiment out of two. (**B)** Competition assay. (Top, left) Scheme of the competition assay. Following a baseline step, the sACE2-Fc (5 µg/ml) was immobilized onto the AHC biosensors. After a second baseline step, a first VHH was applied (VHH1; 5 µg/ml). The sensor was dipped in a mixture of VHH1 and the competitor VHH2 at the same concentration (5 µg/ml). An anti-IgM specific VHH was used as control (ctl). (Top right) Competition assay with immobilized sACE2 pre-bound to B07 (VHH1) and VHH2 being B10, B09 or IgM (ctl). (Bottom left) Competition assay with immobilized sACE2 pre-bound to B09 (VHH1) and VHH2 being B07, B10 or IgM (ctl). (Bottom right) Competition assay with immobilized sACE2 pre-bound to B10 (VHH1) and VHH2 being B07, B09 or IgM (ctl).

To obtain insights into the epitopes recognized by the three VHHs, we carried out a binding competition assay using the BLI system. B07 or B09 did not bind the immobilized ACE2 pre-bound to B07 (ACE2-B07) or B09 (ACE2-B09) (**Fig. 1B**), suggesting that these two VHHs recognize the same, or an overlapping epitope on ACE2. On the contrary, B10 bound ACE2-B07 or ACE2-B09, suggesting that this VHH recognize a different epitope than B07 and B09 (**Fig. 1B**). In agreement with this finding, the immobilized ACE2 pre-bound to B10 (ACE2-B10) was still able to bind B07 or B09 (**Fig. 1B**).

### VHHs B07, B09, and B10 bind ACE2 expressing cells

We tested the ability of the three VHHs to bind ACE2 present at the surface of different cell lines, which express exogenous (HEK293-ACE2, A549-ACE2) or endogenous ACE2 (IGROV-1, Vero E6) (**Fig. 2, Supplementary Fig.1, A and B**). We incubated 10 µg/ml (667 nM) of each VHH with the different cell lines or with parental cells, and we used an unrelated VHH as negative control (ctl). The myc-tagged VHHs were detected bound to the cells’ surface by flow cytometry after staining with an anti-myc antibody and a fluorescent secondary antibody. The three VHHs were specifically detected at the surface of all cell lines, revealing their interaction with trans-membrane ACE2. B10 showed the highest staining in accordance with its high affinity for sACE2 defined by BLI (K_D_ 10 nM) (**Fig. 2A-C, Supplementary Fig. 1, A and B**).

**Fig. 2:**
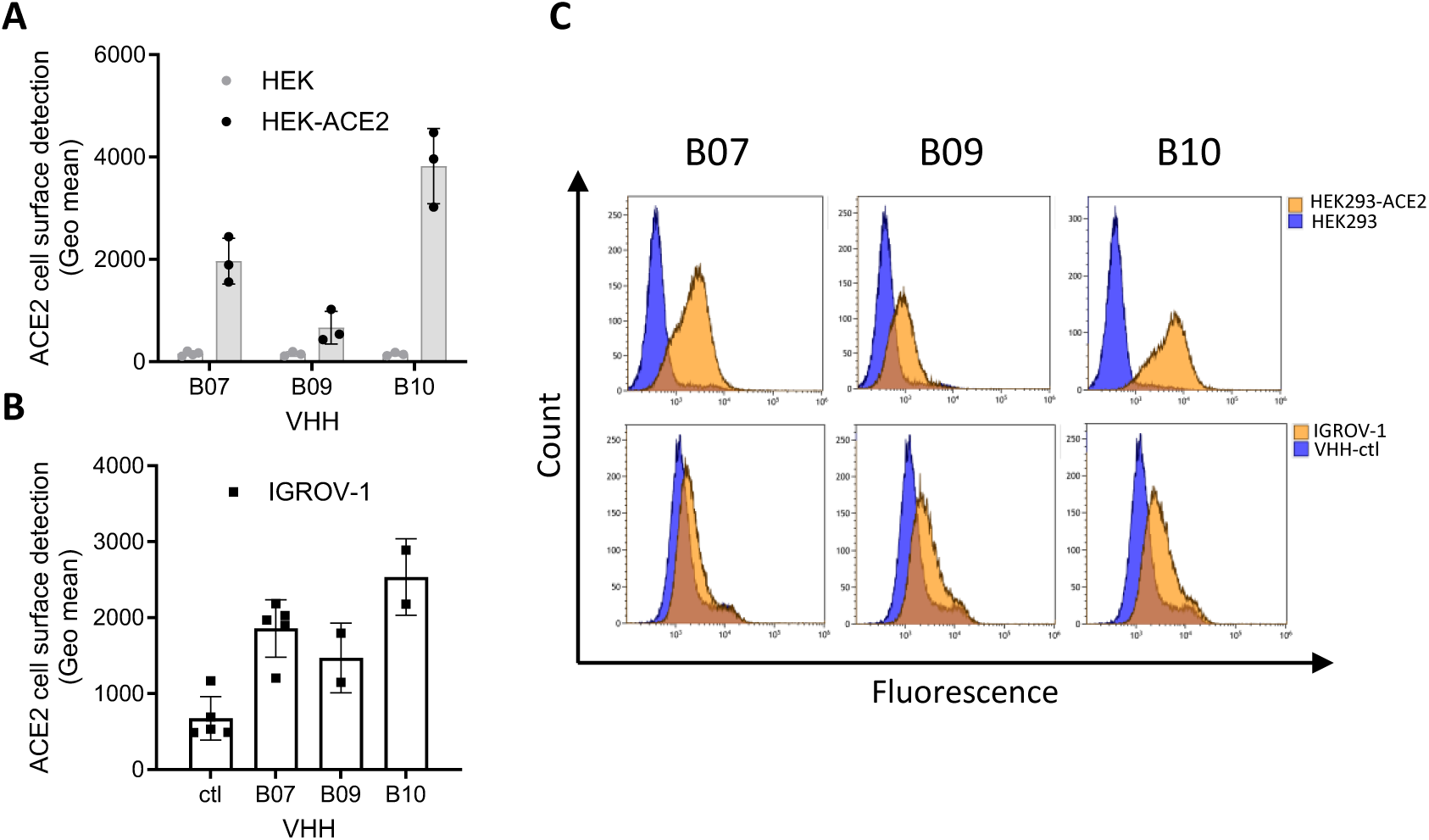
VHHs binding on cells expressing ACE2. Cells were incubated with the different VHHs (10 µg/ml), stained with an anti-myc antibody and a AF488-conjugated anti-mouse antibody, before being analyzed by flow cytometry. (**A)** B07, B09, B10 binding efficacy on HEK293-ACE2-expressing cells. Data are means ± SD of three independent experiments. Parental HEK293 cells were used as control. (**B)** B07, B09, B10 binding efficacy on cells expressing endogenous ACE2 (IGROV-1). Anti-IgE VHH was used as a control (ctl). Data are means ± SD of at least two independent experiments. (**C)** Fluorescence diagram overlays: B07, B09, B10 efficacy on cells expressing exogenous (HEK293-ACE2) or endogenous ACE2 (IGROV-1). Background (blue) corresponds to the fluorescence intensity obtained on parental cells (HEK293) or using a VHH control (anti-IgE VHH) (IGROV-1).

### VHHs B07, B09, and B10 do not impact the enzymatic activity of ACE2

We determined whether B07, B09, and B10 VHHs affected ACE2 function (**Fig. 3**). ACE2 catalyzes the cleavage of small circulating peptides, thus playing a central role in the control of the renin-angiotensin system ^12^. We assayed the impact of 10 µM of each VHH (**Fig. 3A**), or different doses (**Fig. 3C**), on the ability of sACE2 to cleave a fluorogenic peptide substrate (SensoLyte assay). Enzymatic cleavage leads the peptide to fluoresce, which is monitored at excitation/emission wavelength = 330/390 nm. We found no effect of the three VHHs on peptide cleavage (**Fig. 3A**), even at 40 µM of VHHs (**Fig. 3C**), contrary to the inhibitor control DX600 (IC50 = 0.54 µM) (**Fig. 3B**). We confirmed this result *in cellulo* by following peptide cleavage using HEK293-ACE2 live cells instead of soluble ACE2 (**Fig. 3D**). This result suggested that the interaction of B07, B09, and B10 VHHs with sACE2, or cell surface ACE2, did not interfere with its enzymatic activity.

**Fig. 3:**
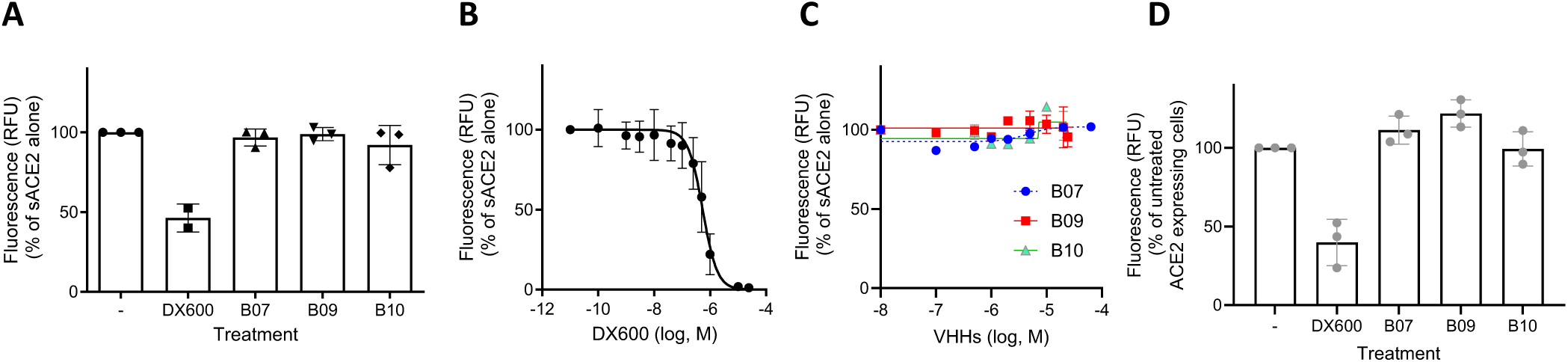
Enzymatic activity of soluble and membrane ACE2. ACE2 activity in the presence of VHHs B07, B09, B10 or absence (−) was assayed using the SensoLyte 390 ACE2 Activity Assay Kit, which measures fluorogenic peptide cleavage. (**A)** Soluble ACE2 (sACE2) activity in the presence of 10 µM VHHs. An ACE2 inhibitor, provided with the kit, was used as control (DX600, 1µM). Data are means ± SD of three independent experiments. (**B)** The ACE2 inhibitor DX600 was used at different doses. Data are means ± SD of seven independent experiments. (**C)** VHHs B07, B09, B10 were used at different doses (one representative experiment out of three performed in duplicates). (**D)** Cell surface ACE2 activity in the presence of saturated concentration of VHHs (> 20 µM) or 1 µM DX600. Data are means ± SD of three independent experiments.

### VHHs B07 and B09 neutralize different SARS-CoV-2 variants with a competitive mechanism

We examined whether the selected VHHs may inhibit the ability of SARS-CoV-2 infected cells to form syncytia using the S-Fuse assay ^30^. We pre-incubated S-Fuse cells (U2OS-ACE2 GFP1-10 and GFP11) with serial dilutions of VHHs, exposed them to different SARS-CoV-2 variants (Delta, BA.1, BQ.1.1, XBB.1.5, XBB.1.16.1, EG.5.1.3; BA.2.86.1) and quantified syncytia formation 18 h after infection.

B07 and B09, which are closely related, inhibited syncytia formation induced with the 7 SARS-CoV-2 variants tested with similar IC50 (from 1.45 to 30 µg/ml – 96.7 to 1970 nM) (**Fig. 4A**). B10 displayed no inhibition activities on the different variants tested (**Fig. 4A**), while having the highest ACE2 affinity. This result was in accordance with the fact that B10 targeted a different epitope than B07 and B09, probably not involved in the interaction between S and ACE2.

**Fig. 4:**
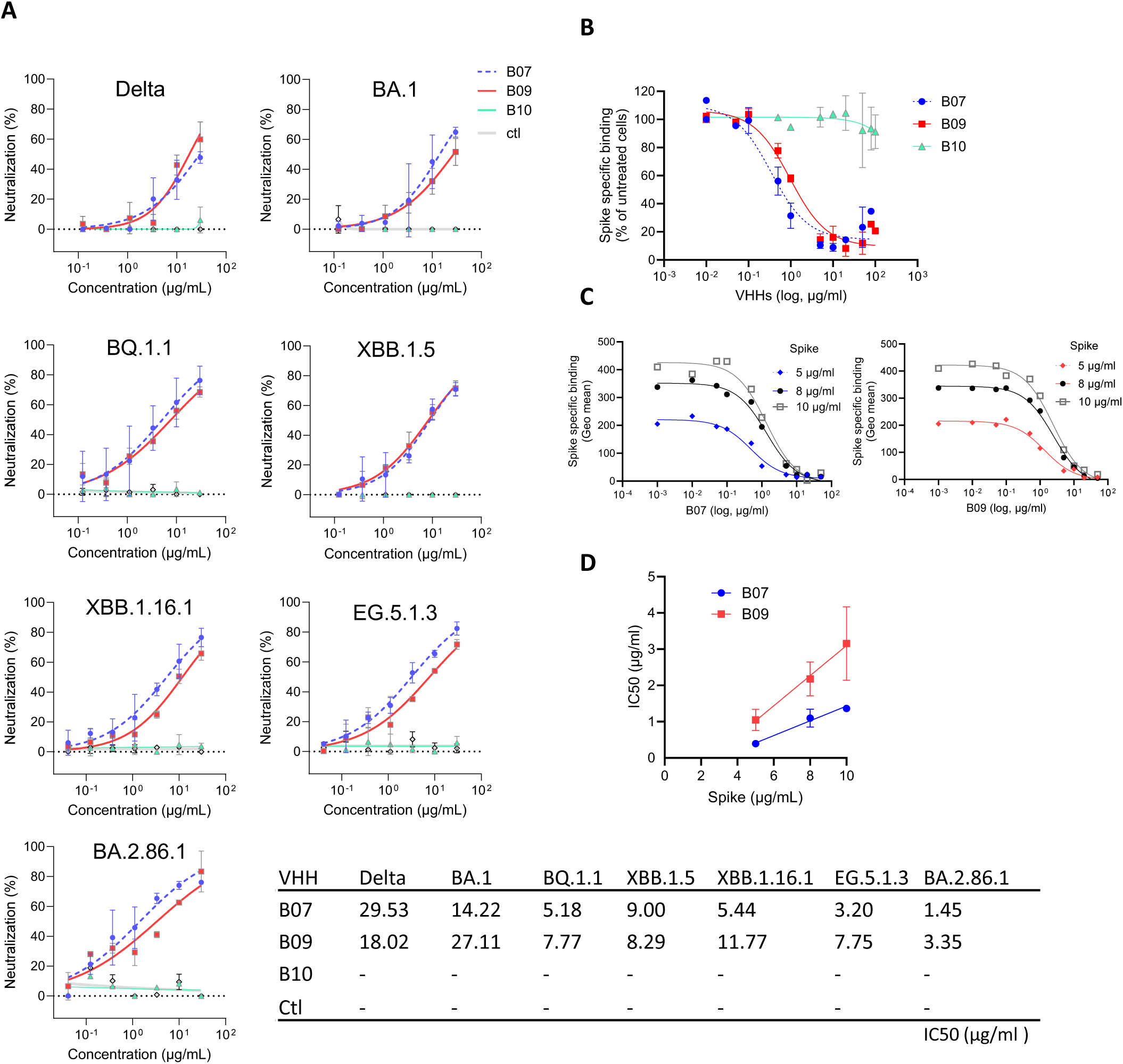
Inhibition of fusion. (**A)** S-fuse assay. Neutralization by B07, B09, B10 VHHs after infection of S-Fuse cells (U2OS-ACE2 GFP1-10 and GFP11) with different SARS-CoV-2 variants (Delta B.1.617.2, BA.1, BQ.1.1, XBB.1.5, XBB1.16.1, EG.5.1.3, BA.2.86.1). Data are mean ± SD of two independent experiments. The dashed line indicates the limit of detection. An anti-IgM (or IgE) specific VHH was used as a control (ctl). IC50 values are indicated in the table. (**B)** Inhibition of 5 µg/ml spike binding to ACE2-expressing HEK293 cells by increasing concentrations of VHHs B07, B09 or B10. HEK293-ACE2 cells pretreated or not with the VHHs and incubated with soluble spike (S) protein (Wuhan) were stained with an anti-S antibody. Results were normalized for nonspecific (0%) and specific binding in the absence of inhibitor (100%). Experiments were fitted to a one-site competitive binding model. Data are mean ± SD of at least two independent experiments. (**C)** Displacement of S binding by B07 (left) or B09 (right) for various concentrations of S (5 µg/ml, 8 µg/ml, 10 µg/ml). Experiments were carried out as in **B** (one representative experiment out of at least two). (**D)** Relationship between the observed IC50 values for B07 or B09 displacement of S binding and initial concentration of S. Results represent means ± SD of at least two independent experiments. Linear regression analysis: R^2^ B07=0.9005; R^2^ B09=0.7896.

To assess whether B07 and B09 neutralized SARS-CoV-2 infection by inhibiting binding to the spike (S) protein, we performed competition experiments in which binding of soluble S (Wuhan) on HEK293-ACE2 cells was measured in the presence of increasing concentration of VHHs B07 and B09. B10 was used as control. S binding was assessed after immuno-staining with an anti-S antibody and flow cytometry (**Fig. 4B**). Pre-incubation of HEK293-ACE2 with VHHs B07 and B09 prevented soluble S binding, while B10 had no effect (**Fig. 4B**). This suggested that B07 and B09 neutralized SARS-CoV-2 infection by inhibiting ACE2-spike interaction.

We then performed competition experiments with B07 or B09 using increasing concentrations of S from 5 to 10 µg/ml (**Fig. 4C**). The relationship between the observed IC50 value and tracer concentration (here S) in displacement experiments indicates whether binding inhibition is competitive (i.e. displaceable) or noncompetitive^31^. The IC50 value is expected to increase linearly with S concentration for competitive inhibition^31^. IC50 values yielded a linear regression with increasing concentration of S (**Fig. 4D**), indicating that both B07 and B09 inhibited S binding through a competitive mechanism.

### VHH-B07 dimerization increases its neutralization efficacy

In order to stabilize the VHHs and increase their efficacy ^25^, we produced dimeric VHHs by fusing the DNA sequence coding for VHH B07 to that of the human IgG1 Fc portion (i.e. hinge plus CH2 and CH3 domains)^32^. We tested the resulting B07-Fc for its ability to interact with sACE2 *in vitro*, to bind membrane ACE2 using different cell types, to prevent the enzymatic activity of ACE2 *in vitro,* and to neutralize SARS-CoV-2 variants (**Fig. 5**).

**Fig. 5:**
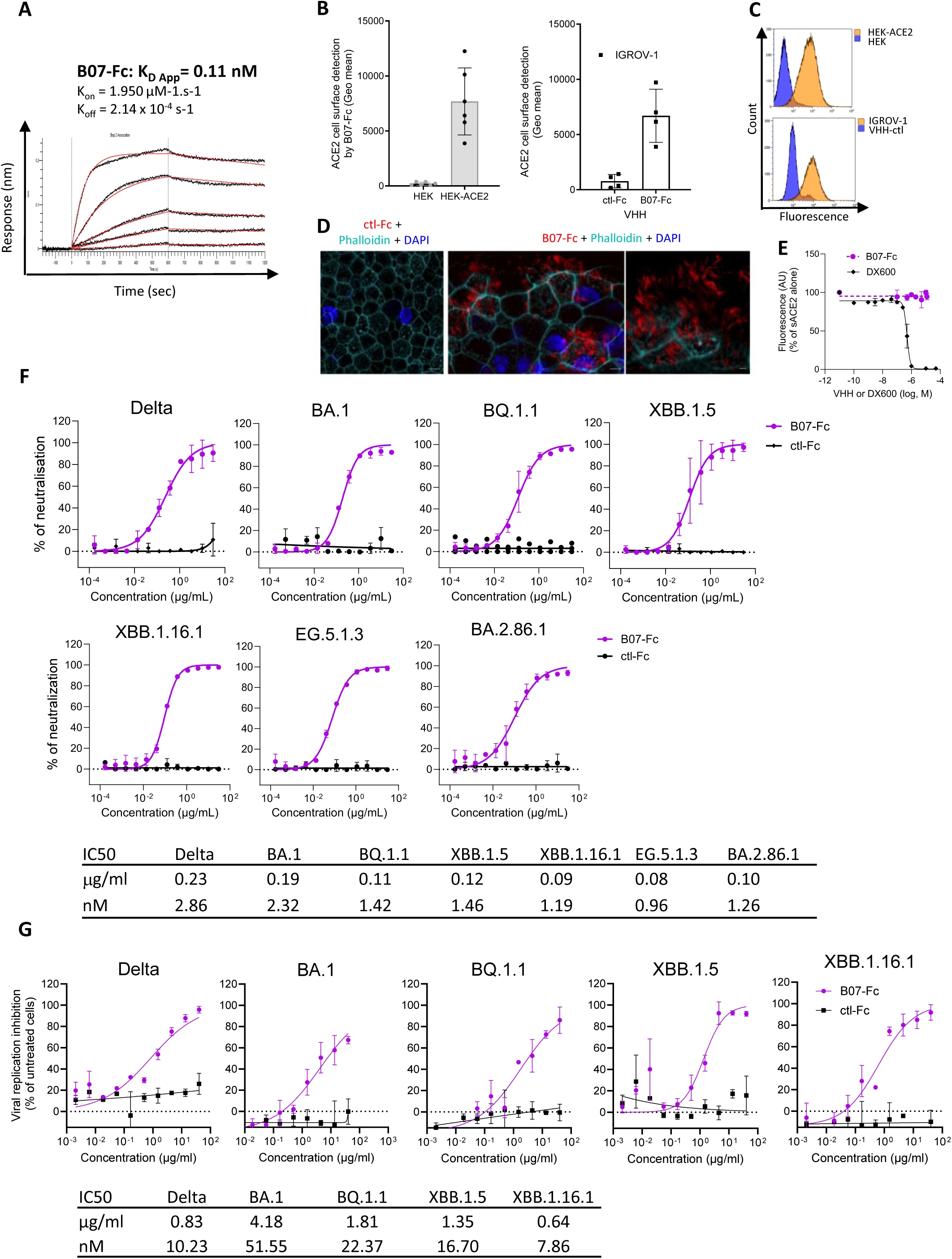
Dimeric VHH-Fc activity. **(A)** Kinetic analysis of VHH B07-Fc on ACE2-His by BioLayer Interferometry (BLI) using various concentrations of VHH B07-Fc (from 0.08 nM to 62 nM). HIS2 biosensors were used to immobilize 1 µg/ml of ACE2-His. (**B)** VHH B07-Fc binding on cells expressing exogenous (HEK293-ACE2) or endogenous ACE2 (IGROV-1). Experiments were performed as in Fig. 2 but using 0.1 µg/ml of VHH B07-Fc. Data are mean ± SD of at least four independent experiments. **(C)** Fluorescence diagram overlays as performed in Fig. 2C. (**D)** ACE2, phalloidin (F-actin) and DAPI staining on primary human nasal epithelial cells (hNECs). Representative immunofluorescence staining of ACE2 (Red: B07-Fc staining) in combination with phalloidin (Cyan) and DAPI (Blue). Scale bars: ctl-Fc, 10 µm; B07-Fc, 5 µm. **(E)** Enzymatic activity of soluble ACE2 (sACE2) in the presence of VHH B07-Fc or absence using the SensoLyte 390 ACE2 Activity Assay Kit, as performed in Fig. 3. DX600 was an inhibitor control of the kit (one representative experiment out of two). **(F)** Inhibition of fusion (S-Fuse assay) by B07-Fc after infection of S-Fuse cells with different SARS-CoV-2 variants : Delta B.1.617.2, BA.1, BQ.1.1, XBB.1.5, XBB.1.16.1, EG.5.1.3, BA.2.86.1. The dashed line indicates the limit of detection. Data are mean ± SD of at least two independent experiments. **(G)** Inhibition of SARS-CoV-2 replication by B07-Fc measured by RT-qPCR. Vero E6 cells pre-incubated with B07-Fc were infected with Delta B.1.617.2, BA.1, BQ.1.1, XBB.1.5 and XBB.1.16.1 (MOI between 0.07 and 0.33) during 48 or 66 hours and viral replication was measured by quantitative RT-qPCR (one experiment in triplicates).

B07-Fc bound sACE2 with higher avidity (x3,500) compared to monomeric B07 (**Fig. 1**), with a measured apparent K_D_ (K_D_ app) of 0.11 nM (**Fig. 5A**). B07-Fc was specifically detected at the surface of different cell lines expressing exogenous (HEK293-ACE2, A549-ACE2) or endogenous ACE2 (IGROV-1, Vero E6) by flow cytometry after staining with anti-IgG antibodies (**Fig. 5, B and C, Supplementary Fig. 1, C and D**). B07-Fc also specifically stained primary human nasal epithelial cells (hNECs), as shown by confocal immunofluorescence imaging (**Fig. 5D**). This air-liquid culture system is an effective tool for studying SARS-CoV-2 infection^33^. As described^34,35^, ACE2 co-localized with α-tubulin on the surface of motile cilia (**Supplementary Fig. 2A**). As seen with B07, B07-Fc did not affect enzymatic peptide cleavage activity of sACE2 (**Fig. 5E**) and did not impact cell viability, even at high concentration (**Supplementary Fig. 2B**). Finally, B07-Fc inhibited syncytia formation induced with the 7 SARS-CoV-2 variants and sub-variants tested (Delta, BA.1, BQ.1.1, XBB.1.5, XBB.1.16.1, EG.5.1.3, BA.2.86.1)^9^ (**Fig. 5F**) with an IC50 around 2 log lower than that of B07 (from 0.08 to 0.23 µg/ml-0.96 to 2.86 nM). B07-Fc also inhibited infection of Delta and Omicron variants (BA.1, BQ.1.1, XBB.1.5, XBB.1.16.1) with IC50 from 0.6 to 4.2 µg/ml (7.9 to 22.4 nM) (**Fig. 5G**). These results suggested that the dimerization of the VHH B07 increased its neutralizing efficacy without affecting its enzymatic activity. Therefore, such a VHH represented a good candidate for further *in vivo* studies.

### ACE2 N-terminal variation impacts VHH B07-Fc binding efficacy

We evaluated whether human ACE2 (hACE2) polymorphism affect B07-Fc binding efficacy. We constructed polymorphic hACE2 variants by substituting residues located within the vicinity of S binding (S19P, I21T, K26R, T27A, H34N, E35K, E37K, K68E, M82I, P84T) in a myc-hACE2 expressing plasmid. We tested their cell surface expression after anti-myc staining and their reactivity with B07-Fc by flow cytometry. None of the substitution altered the correct cell surface expression of myc-hACE2 constructs (**Supplementary Fig. 3A**). Eight of ten constructs were correctly detected by B07-Fc, suggesting that hACE2 polymorphism might have little impact on B07-Fc antiviral efficacy (**Fig. 6A**). However, B07-Fc failed to detect hACE2-H34N and hACE2-E35K expressing cells (**Fig. 6A**), revealing the importance of H34 and E35 for B07-Fc binding to hACE2. The substitution of E35 by an Alanine (E35A) abrogated B07-Fc binding, confirming that E35 may be part of the epitope (**Supplementary Fig. 3B**).

**Fig. 6:**
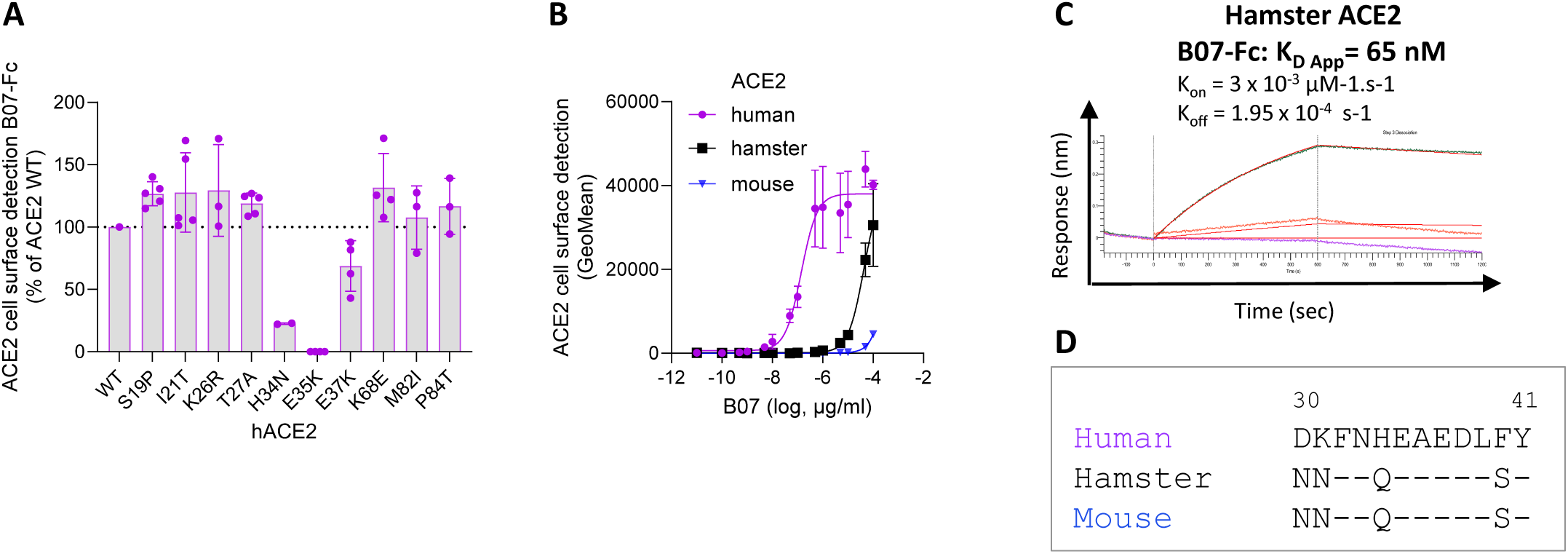
Impact of hACE2 variation on VHH B07-Fc binding. (A, **B)** Cells were transfected with different ACE2 constructs, incubated with B07-Fc (0.1 µg/ml or different doses) and stained with an AF647-conjugated anti-human antibody before being analyzed by flow cytometry. Data are mean ± SD of at least two independent experiments. **A)** Cells transfected with hACE2 polymorphism constructs. (**B**) Cells transfected with human, hamster, or mouse ACE2. (**C**) Kinetic analysis of B07-Fc on hamster ACE2-His by BioLayer Interferometry (BLI) using various concentrations of VHH B07-Fc (from 18 to 595 nM). HIS2 biosensors were used to immobilize 2 µg/ml of His-ACE2. K_D_ are indicated. **(D)** Alignment of the amino acid sequences of human, hamster, and mouse ACE2 from position 30 to 41.

Phylogenetic diversities in ACE2 might also influence B07-Fc binding. B07-Fc poorly recognized hamster and mouse ACE2 expressing cells compared to hACE2 expressing cells (**Fig. 6B**). BLI experiments confirmed that B07-Fc bound soluble hamster ACE2 with smaller avidity (3 log) compared to human ACE2 (**Fig. 6C**, **Fig. 5A**). The presence of a Glutamine at position 34 (Q34) in these different species **(Fig. 6D**) may explain this lower binding efficacy. Indeed, B07-Fc did not react with hACE2-H34Q (**Supplementary Fig. 3B**). Therefore, the B07-Fc epitope seems critically dependent upon residues H34 and E35 of hACE2.

### Fc-conjugated B07 dimers protect against Omicron infection *in vivo*

We evaluated *in vivo* the prophylactic potential of neutralizing VHH B07-Fc using hACE2 transgenic mouse model (K18-hACE2) and XBB.1.5 Omicron variant infection. Mice treated intranasally with a single administration of VHH B07-Fc (7 mg/kg), or VHH-Fc control (7 mg/kg) were infected 24 h later intranasally with 1×10^5^ PFU of XBB.1.5 variant. Animals were euthanized at day 3 post-infection and lung viral load was measured by RT-qPCR (**Fig. 7A**). In 6 of the 10 B07-Fc-treated mice, lung viral load level was strongly reduced (> 3 log) or below the detection limit, suggesting that the lung had been protected from viral infection (**Fig. 7B**). We assessed by immunofluorescence the cytopathic effect induced by XBB.1.5 in the presence (B07-Fc) or absence (ctl-Fc) of B07-Fc. Nasal, respiratory and lung epithelia were stained at day 3 p.i. with phalloidin (to visualize F-actin), anti-alpha tubulin antibodies (to visualize cilia) and Dapi (to visualize the nucleus). Protected mice showed no tissue damage (disappearance of the ciliated structure) compared to control mice (ctl-Fc and uninfected K18-hACE2) (**Fig. 7C**), corroborating the protector effect of B07-Fc. The dose of B07-Fc in the lung homogenate measured by ELISA was variable between treated mice, revealing probable heterogeneity in the diffusion of VHH in the lung. Notably, all mice in which the B07-Fc dose in the lungs was above 215 ng B07-Fc of lung extract had lung viral load below the limit of detection, while the mice with lower B07-Fc dose exhibited high lung viral load (**Fig. 7D**). Therefore, we concluded that B07-Fc, when used prophylactically in K18-hACE2 and when efficiently delivered into the lungs, strongly protects against Omicron pulmonary infection.

**Fig. 7:**
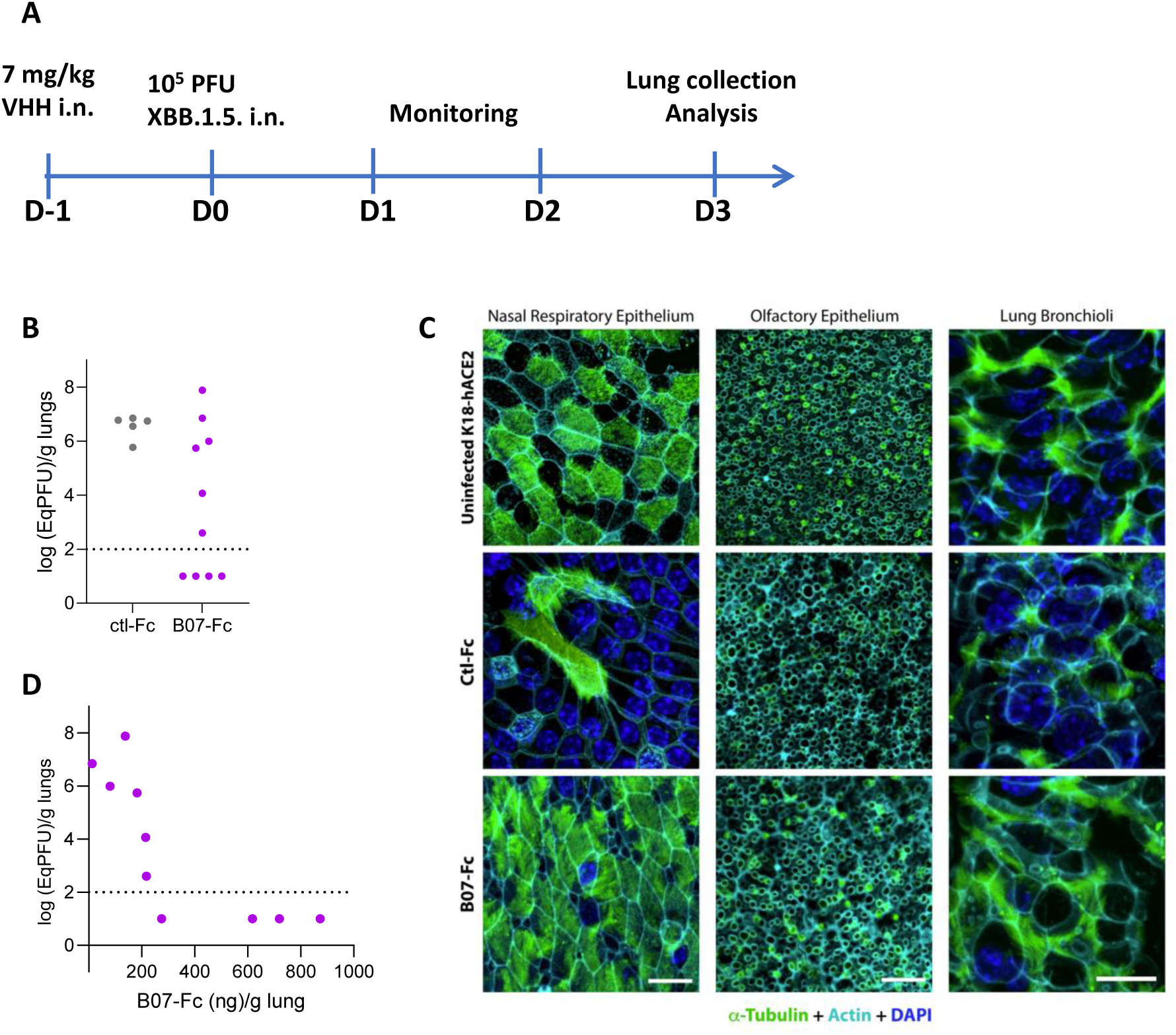
Prophylaxis of VHH B07-Fc for SARS-CoV-2 infection in mice. **(A)** Schematic diagram showing the experimental design of B07-Fc prophylaxy in XBB.1.5 infected K18-hACE2 mice. Animals received intranasally (i.n.) 7 mg/kg VHH-B07-Fc (B07-Fc) or 7 mg/ml VHH-Fc ctl (ctl-Fc). Twenty four hours later, they were infected with 10^5^ PFU XBB.1.5. intranasally (i.n.). Three days post-infection, lungs were collected for analysis. **(B)** RNA load measured by RT-qPCR of SARS-CoV-2 in lung. **(C)** Tubulin, phalloidin (F-actin) and DAPI staining on nasal respiratory epithelium, olfactory epithelium and lung bronchioli extracted from K18-hACE2 uninfected mice, and animals that were received VHH-Fc (Ctl-Fc or B07-Fc). Representative immunofluorescence staining of Tubulin (Green) in combination with phalloidin (Cyan) and DAPI (Blue). Scale bars: 10 µm. **(D)** Relationship between RNA load measured in the lungs by RT-qPCR and B07-Fc dose quantified in the lungs by ELISA. ELISA plates coated with sACE2 and incubated with homogeneate of lower lung lobes of B07-Fc treated animals or a range of purified B07-Fc as a control were stained with peroxidase anti-human IgG.

Next, we evaluated the prophylactic potential of VHH-B07-Fc using the Syrian hamster preclinical model. VHH-B07-Fc was administered intranasally at 5 mg/kg and infected 24h later with 1×10^4^ PFU of the XBB.1.16.1 variant. Lung viral load was modestly but significantly lower in hamster treated with B07-Fc compared with the control animals (3.49×10^7^ vs 5.7×10^8^ copies/g lung) (**Supplementary Fig. 4**), indicating a lower protection potential of VHH-B07-Fc in this species compared with transgenic mice expressing human ACE2.

## Discussion

Targeting ACE2 represents an alternative to the use of mAbs targeting the spike (S) protein. Indeed, the anti-ACE2 VHHs conferred neutralization against a broad spectrum of variants including Omicron subvariants XBB.1.16.1, EG.5.1.3 and BA.2.86.1 (**Fig. 4 and Fig. 5**). In comparison, the therapeutic mAb sotrovimab weakly neutralizes XBB.1.16.1 and EG.5.1.3 and lost antiviral activity against BA.2.86.1 and JN.1^36^ ^9^. This underlines the power of this host targeting approach.

An issue of host targeting is the potential loss of physiological functions of the target. The anti-ACE2 VHHs bind the peptidase domain of ACE2 (**Fig. 1**) without interfering with its peptidase activity *in vitro* and *in cellulo* (**Fig. 3**, **Fig. 5E**). They neutralized SARS-CoV-2 infection by competitively inhibiting the interaction with S (**Fig. 4B-D**), targeting an epitope comprising residues H34 and E35 of the N-terminal helix 1 of ACE2 (**Fig. 6A**). This epitope, distal to the enzyme active site (residues 340-450), should not induce side effects if it is targeted. This suggests the translational potential of these antibodies. Moreover, the emergence of escape mutants after treatment with competitive anti-ACE2 VHHs is unlikely, underscoring the value of this host-directed therapy. Resistance would require a change in the way S binds to the ACE2/VHH complex, likely incompatible with the fusogenic activity of the protein.

Compared with conventional mAb-based therapies, VHH-based therapies have several advantages due to their small size and their excellent solubility and stability across a wide temperature range^38^. A particular asset of VHHs is the possibility of being administrated by inhalation, which is a non-invasive method of delivery^39^. Consistent with this, B07-Fc prevented the infection of human ACE2-expressing mice (K18-hACE2) by Omicron XBB.1.5 following intranasal administration (**Fig. 7**). The observed heterogeneity in the number of fully protected mice correlated with the effective concentration of B07-Fc retrieved in the lungs, likely reflecting a sub-optimal distribution of B07-Fc, although VHH degradation or aggregation cannot be completely excluded. Beyond a threshold dose of B07-Fc in the lung (215 ng of B07-Fc/g lung), no viral load was detected in animals. We confirmed this protective effect of B07-Fc after intranasal administration in the preclinical Syrian hamster model^40–42^ (**Supplementary Fig. 4**), although this protection was reduced. The weak binding of B07-Fc to hamster ACE2 compared to hACE2 (**Fig. 6C**) likely explained the low effectiveness of B07-Fc in hamster. This observed protection of animals suggests that without any additional development in the formulation, VHH B07-Fc is already highly stable to be used by inhalation. However, the administration procedures require optimization to further boost overall protection. Intranasal administration of VHHs already proved its efficacy to prevent SARS-CoV-2 infection in animal models^43–45^ and to reduce the viral load in the brain^46^. In the case of anti-ACE2, this mode of administration, by promoting bio-distribution in the lung, may also overcome a loss of anti-ACE2 activity due to the wide expression of ACE2 in different tissues and the presence of circulating ACE2 in the blood^13^. This would result in the need of much reduced dose to achieve antiviral activity. In addition, respiratory drug delivery presents low logistical burden and possible self-administration, which would be particularly beneficial in the case of SARS-CoV-2^43^.

Another advantage of VHH is that they are simple to engineer. The availability of VHH DNA sequences allows rapid modification of antibody fragments. Engrafting B07 onto the Fc domain of conventional IgG1 results per se in a dimeric VHH, with an increased avidity and neutralization efficacy, which rivals those of anti-ACE2 mAbs^20–22^ (**Fig. 5, F and G**), while showing no toxicity on cells (**Supplementary Fig. 2B**). In order to further enhance activity, VHHs could be engineered to combine two or three VHHs in a single polypeptide to target distinct epitopes or even distinct receptors^47^. This would lead to synergistic effects^48^. The fusion with conventional Fc domain allows VHHs to bind Fc receptors, thereby rescuing them from degradation, improving their pharmacokinetic profile^25,26^ and increasing their half-life *in vivo*^27^. However, fusing a VHH to an Fc fragment, may activate Fc effector function on immune cells or with complement, or cause antibody-dependence enhancement (ADE)^49^. To mitigate these risks and increase the potential of therapeutic properties, optimal balance of Fc effector functions can be modulated by the introduction of point mutations in the Fc domain^50^ ^51^. It could be of interest to explore the effect of these known mutations on the stability and efficacy of VHH B07-Fc.

Beside their inhibitor activity against SARS-CoV-2, VHHs against ACE2 represent useful tools to answer fundamental questions regarding the SARS-CoV-2 entry process. VHHs B10 and B07-Fc bind ACE2 with high efficacy (K_D_ app. ≤ 10 nM) and may allow the detection of ACE2 in cryptic locations in primary cells not accessible to conventional antibodies (**Fig. 5D**). B10, which is not neutralizing, represents a promising tool to track ACE2 in SARS-CoV-2 infected cells. Anti-ACE2 VHH also offers the possibility to study the soluble circulating form of ACE2^13^ and its relationship with viral, kidney or heart diseases^52^. The concentration of sACE2 may correlate with systemic inflammation^53^ or myocardial injury^54^, but its role is largely unknown.

## Methods

### Expression and purification of ACE2 peptidase domain

For alpaca immunization the ACE2 peptidase domain (residues 19-615, Uniprot: Q9BYF1) was cloned in pcDNA3.1. The signal peptide from the IgK chain (METDTLLLWVLLLWVPGSTG) was introduced at the N-terminus to ensure the protein secretion to the extracellular medium, and His(x8) and Strep tags were added *in-tandem* at the C-terminus, following a Thrombin cleavage site.

The plasmid was transiently transfected into Expi293F^TM^ cells (Thermo-Fischer) using FectoPro^®^ DNA transfection reagent (PolyPlus) according to the manufacturer’s instructions, and 5 μM kifunensine was added to the culture medium immediately after transfection. The cells were incubated 5 days at 37 °C and harvested by centrifugation. The supernatant was concentrated and recombinant ACE2 was purified by affinity chromatography on a Streptactin column (IBA) and by size-exclusion chromatography (SEC) on a Superdex200 column (Cytiva) equilibrated 10 mM Tris, 100 mM NaCl. ACE2 was digested over-night at room temperature with Thrombin (1 U per 100 μg of recombinant protein) to remove the His and Strep tags. The proteolysis was stopped with 1 mM PMSF (phenylmethylsulfonyl fluoride) and the untagged ACE2 was recovered in the flow-through after injection in a Streptactin column. A final SEC was performed, after which untagged ACE2 was concentrated and stored at −80 °C. For competition experiments using BLI, ACE2 peptidase domain fused to a human Fc from IgG1 ^32^ was used (sACE2-Fc). For the measurement of the affinity between VHH B07-Fc and sACE2 by interferometry (BLI), we used a His-tagged sACE2 (sACE2-His) that was expressed as indicated above and purified by IMAC and size-exclusion chromatography.

### Plasmids and transfection

Mouse ACE2 plasmids was purchased from addgene (pscALPSpuro-MmACE2, #158087). Hamster ACE2 pcDNA3.1 hygro plasmid was synthetized by GeneArt (ThermoFisher). The construct human ACE2 (hACE2) was obtained by substitution of the sequence encoding mCardinal from mCardinal-C1 plasmid (addgene, #54799) with hACE2 from the pLenti6-attB-hACE2-BSD^30^. The construct myc-hACE2 (myc-hACE2) was obtained by substitution of the sequence encoding mCardinal from mCardinal-C1 plasmid (addgene, #54799) with myc-hACE2 from the pCEP4-myc-ACE2 (addgene, #141185)^55^. Polymorphism substitutions were generated by site-directed mutagenesis using the Q5-site directed mutagenesis kit (New England Biolabs) according to the manufacturer’s instructions. The mutants were all verified by sequencing (Eurofins). For transient expression, HEK293 (2×10^5^) cells were cotransfected with WT or mutant ACE2 vectors and with EGFP-N1 (Clontech, Palo Alto, Calif.), in a 5:1 ratio. The EGFP positive cells were analyzed by flow cytometry after immunostaining.

### Viruses

SARS-CoV-2 delta (B.1.617.2; GISAID ID: EPI_ISL_2029113), and omicron BA.1 (GISAID ID: EPI_ISL_6794907), and BQ.1.1 (GISAID ID: EPI_ISL_15731523) were previously described ^36,56–58^. XBB.1.5 was isolated from a nasopharyngeal swab of an individual of Hôpital Européen Georges Pompidou (HEGP; Assistance Publique, Hôpitaux de Paris) (GISAID ID: EPI_ISL_16353849)^59^. The laboratory of Virology of HEGP sequenced the swabs. The patient provided informed consent for the use of the biological materials. The XBB.1.16.1 strain (hCoV-19/France/GES-IPP07712/2023), the EG.5.1.3 strain (hCoV-19/France/BRE-IPP15906/2023), and the BA.2.86.1 strain (hCoV-19/France/IDF-IPP17625/2023) were supplied by the National Reference Centre for Respiratory Viruses hosted by Institut Pasteur (Paris, France) and headed by Dr Etienne Simon-Lorière^9^. The human sample from which strain hCoV-19/France/GES-IPP07712/2023 was isolated has been provided by Dr Vanessa COCQUERELLE from Laboratory Deux Rives. The human sample from which strain hCoV-19/France/BRE-IPP15906/2023 was isolated has been provided by Dr F. Kerdavid from Laboratoire Alliance Anabio, Melesse. The human sample from which hCoV-19/France/IDF-IPP17625/2023 was isolated was provided by Dr Aude LESENNE from Cerballiance, Lisses (France). Titration of viral stocks was performed on Vero E6 or IGROV-1 cells, with a limiting dilution technique enabling the calculation of the median tissue culture infectious dose or on S-Fuse cells.

In replication experiments, XBB.1.16.1 (hCoV-19/France/GES-IPP07712/2023), XBB.1.5 (hCoV-19/France/PDL-IPP58867/2022), BQ.1.1 (hCoV-19/France/IDF-IPP50823/2022) and BA.1 (hCoV-19/France/PDL-IPP46934i/2021) were supplied by the National Reference Centre for Respiratory Viruses hosted by Institut Pasteur (Paris, France). The human samples from which strains were isolated has been provided from the Laboratoire Deux Rives, CH Laval and Laboratoire des centres de santé et d’hôpitaux d’IDF. Viral productions were performed on Vero-TMPRSS2 cells and titration of viral stocks was performed on Vero E6 cells. Viral concentrations were determined through Plaque Assay and the titers were quantified as plaque forming units (pfu) per milliliter.

The sequence of the viral stocks was verified by RNAseq by the National Reference Centre for Respiratory Viruses hosted by Institut Pasteur (Paris, France). All work with infectious virus was performed in biosafety level 3 containment laboratories at Institut Pasteur.

### Cell lines

HEK293-ACE2 (human embryonic kidney cell line) were generated by transfection and cultured for several weeks in Blasticidin (10 µg/ml). U2OS (human osteosarcoma cell line) and A549 (human lung epithelial cell line) cells stably expressing ACE2, and U2OS-ACE2 cells stably expressing the GFP split system (GFP1-10 and GFP11; S-Fuse cells) were previously described ^30^. These cells were cultured in DMEM or in F-12K Nutrient Mixture Media (A549) supplemented with 10% fetal bovine serum (FBS), and 1% penicillin/streptavidin. Blasticidin (10 μg/mL) and puromycin (1 μg/mL) were used to select for ACE2 and GFP split transgenes expression, respectively.

Vero E6 (Vero 76, clone E6, Vero E6, ATCC® CRL-1586TM, simian kidney cell line) was obtained from ATCC (USA) and cultured in DMEM medium (Gibco) containing 10 % (v/v) FBS (Gibco) and penicillin/streptavidin (ThermoFisher Scientific). IGROV-1 cells (Human ovarian cancer cell line) were from the NCI-60 cell line panel ^36^ and cultured in RPMI medium (ThermoFisher) containing 10% FBS and penicillin/streptavidin. Vero E6 and IGROV-1 endogenously express ACE2.

Absence of mycoplasma contamination was confirmed in all cell lines with the Mycoalert Mycoplasma Detection Kit (Lonza). All cell lines were cultured at 37°C and 5 % CO_2_.

### Animal immunization and library construction

All immunization processes were executed according to the French legislation and in compliance with the European Communities Council Directives (2010/63/UE, French Law 2013-118, February 6, 2013). The Animal Experimentation Ethics Committee of Pasteur Institute (CETEA 89) approved this study (2020-27412). We subcutaneously injected an adult alpaca at days 0, 21, and 28 with approximately 150 µg of ACE2 mixed with Freund complete adjuvant for the first immunization and with Freund incomplete adjuvant for the following immunizations. A blood sample of about 150 ml of the immunized animal was collected and Peripheral Blood Mononuclear cells (PBMCs) were isolated on a ficoll discontinuous gradient. Briefly, the *vhh* genes isolated from PBMCs were then cloned into the phagemid vector pHEN6 by using primers contained enzymatic SfiI and NotI restriction sites at the 5’ and 3’ ends, respectively. The size of the library was estimated at about 8×10^7^ cfu. Phage display protocol was performed as described in Lafaye et al. ^60^. 10^12^ phage-VHH diluted in PBS were used to perform three rounds of panning by using sACE2 coated on Nunc Immunotubes (Maxisorp) tubes (10 µg/ml). To increase the stringency, different blocking buffers for each panning were used: 2% skimmed milk as saturating agent for the first round; 5% BSA (Bovine Serum Albumin) and Odyssey blocking buffer (LI-COR Biosciences) diluted at 1/2 for the second and third one.

Following panning, phage clones were screened by standard phage-ELISA procedures using a HRP/anti-M13 monoclonal antibody conjugate. The DNA corresponding to the positive VHHs was sequenced by GATC Biotech. The sequences were processed with DNA strider and analyzed using ClustalW2-Multiple Sequence Alignment of EMBL-EBI.

### VHH production

The pHEN6 vector allows the periplasmic expression and purification of selected VHHs with a c-myc tag and His tag at the C terminal end in frame with the VHH. Transformed *E. coli* TG1 cells expressed VHHs in the periplasm after induction with IPTG (1 mM) for 16h at 30°C. After centrifugation, cells were resuspended in PBS then lysed by polymixin B sulfate (1 mg/ml) for 1h at 4°C, the periplasmic extract being obtained after centrifugation. Purified VHHs were obtained by His pure Cobalt sepharose beads (Thermo) according to the manufacturer’s instructions.

### Dimeric VHH-Fc production

The coding sequences of the selected VHHs were digested with NcoI and NotI and subcloned into pFuse-huIgG-Fc2 digested with NcoI and NotI ^61^. *E. coli* XL1-Blue transformants were obtained on 2YT agar plates containing zeocin 25 µg/ml. The plasmids coding for the recombinant proteins were purified with Nucleobond Xtra Midi Plus EF (Macherey Nagel), transiently transfected in Expi293™ cells (ThermoFisher Scientific) using Fectro PRO DNA transfection reagent (Polyplus), according to the manufacturer’s instructions. Cells were incubated at 37°C for 5 days and then the cultures were centrifuged. Proteins were purified from the supernatants by affinity chromatography using a HiTrap protein A HP (Cytiva), followed by Size Exclusion Chromatography on a Superdex 75 column (Cytiva) equilibrated in PBS. Peaks corresponding to the dimeric VHH-Fc proteins were concentrated and stored at −80°C until used.

### Biolayer interferometry (BLI)

Equilibrium binding was performed via BLI-analysis using an Octet HTX instrument (Octet HTX, Sartorius). VHH-His (or sACE2-His) (5 µg/ml) diluted into Sartorius’s PBS working buffer were immobilized onto Ni-NTA biosensors (Sartorius), which were pre-equilibrated in the same buffer. Following a baseline, VHH-coated sensors (or sACE2 coated sensors) were incubated with various concentrations of sACE2 (0.3; 0.625; 1.25; 2.5; 5; 10; 20 µg/ml) (or various concentration of VHH-B07-Fc: 0.15; 0.3; 0.625; 1.25; 2.5; 5; 10 µg/ml) for 10 min to reach equilibrium. Subsequently, biosensors were put in working buffer for 10 min to initiate dissociation. All incubations were performed at a temperature of 30°C under continuous shaking (1,000 rpm). Data were analyzed using the Octet Software 11.0. Binding curves from the association and dissociation steps were fitted using a 1:1 binding model to determine K_D_.app values.

To compare B07-Fc binding on human versus hamster ACE2, sACE2-His (1 or 2 µg/ml)-coated HIS2 sensors (Sartorius) were incubated with various concentration of VHH-B07-Fc: from 0.08 nM to 62 nM (human ACE2) or from 18 to 595 nM (hamster ACE2).

To measure competitive binding to sACE2 protein between the different VHHs by BLI, we immobilized ACE2 fused to the human Fc domain (sACE2-Fc) onto Anti-hIgG Fc Capture (AHC) Biosensors (Sartorius) at 5 µg/ml for 10 minutes. A baseline step was realized by dipping sensors into the working buffer. Then, a first VHH was applied at 5 µg/ml, in order to bind the immobilized ACE2 and reach saturation. The sensor was then dipped into a well containing a mixture of the same VHH with the competitor one added at the same concentration (5 µg/ml).

### Measurement of VHH binding or spike binding on ACE2 expressing cells by flow cytometry

Cells (2×10^5^) were incubated for 60 min at 4°C with 10 µg/ml VHHs or 0.1 µg/ml VHH-Fc, washed twice with 2% FBS-PBS, incubated for 60 min at 4°C with anti-myc antibody (9E10, Abnova), washed twice with 2% FBS-PBS, and incubated for 60 min at 4°C with AF488-conjugated anti-mouse antibody (Invitrogen) (for VHHs detection), or with the alexa-Fluor-488-conjugated anti-human IgG antibody (Invitrogen) (for VHH-Fc detection). The stained cells were washed twice, fixed in PFA 1.5% and analyzed on Attune flow cytometer NxT flow cytometer (ThermoFisher) and Kaluza software. The parental cells (HEK293, A549) or a VHH control (anti-IgE or anti-IgM VHHs) were used as negative controls.

To perform competition experiments of soluble spike binding an increasing concentration of VHH (between 0.01 to 100 µg/ml) were incubated with HEK293-ACE2 cells in RPMI/10% FBS. After 30 min at 37°C, 5 µg/ml, or different doses (5, 8, 10 µg/ml), of soluble trimeric spike (Wuhan) ^62^ was added for 30 min at 37°C. Then, immuno-staining with 5 µg/ml anti-spike monoclonal antibody mAb48 (gift of Hugo Mouquet, institute Pasteur) ^63^ and alexa-Fluor-488-conjugated anti-human IgG antibody (Invitrogen) was performed at 4°C. Fixed cells (PFA 4%) were analyzed on Attune flow cytometer NxT flow cytometer (Thermo Fisher) and analyzed as described above.

### ACE2 enzymatic activity

Human ACE2 activity was evaluated using SensoLyte 390 ACE2 Activity Assay Kit (AnaSpec) according to the manufacturer’s protocol. The activity was measured on purified sACE2 (10 nM). B07, B09, and B10 were added at different concentrations before activity measurement. The protease activity of ACE2 is indicated by an increase of the fluorescence after peptide cleavage.

To measure ACE2 activity on cells, 2.5×10^4^ HEK293-ACE2 cells per well (96-well black plate, Greiner-BioOne) were incubated with B07, B09, B10 (>20 µM) or the control inhibitor DX600 (1µM) in 40 µl complete medium without phenol red (Gibco) before adding 40 µl of substrat diluted in PBS (SensoLyte 390 ACE2 Activity Assay kit). Peptide cleavage was measure in the supernatant on Tecan SPARK plate reader.

### SARS-CoV-2 neutralization assay: S-Fuse

U2OS-ACE2 GFP1-10 and GFP11 cells, also termed S-Fuse cells, become GFP+ when they are productively infected by SARS-CoV-2 ^30^. Cells were mixed (ratio 1:1) (U2OS-ACE2 GFP1-10 (10^4^) and GFP11 cells (10^4^)) and plated in a μClear 96-well plate (Greiner Bio-One). Serial dilution of VHHs were incubated with the cells at 37°C and infected with a SARS-CoV-2 strain. Eighteen hours later, cells were fixed with 4% paraformaldehyde (PFA), washed and stained with Hoechst (dilution 1:10000, Invitrogen). The extent of fusion was then quantified by measuring the number of objects GFP+ with an Opera Phenix high-content confocal microscope (PerkinElmer) and Harmony software version 4.9 (PerkinElmer). The percentage of inhibition was calculated using the number of syncytia as value with the following formula: 100 × (1 − (value with VHH −value in ‘non-infected’)/(value in ‘control VHH’ − value in ‘non-infected’)). Inhibition activity of each VHH was expressed as the IC50. We previously reported that the neutralization assay with the S-Fuse system is not affected by differences in fusogenicity between variants^64^.

### SARS-CoV-2 neutralization assay: Viral replication

For the evaluation of SARS-CoV-2 replication, the number of SARS-CoV-2 copies was measured by reverse transcription quantitative polymerase chain reaction (RT-qPCR). Individual VHHs were pre-incubated with Vero E6 cells (ten specified concentrations) 2 hours prior infection. Each plate included PBS (2 µl) and remdesivir (25 µM; SelleckChem) controls. After the pre-incubation time, the cells were exposed to the different variants of SARS-CoV-2 (at a multiplicity of infection between 0.07 and 0.33 PFU/Vero E6 cells). After a one-hour adsorption at 37°C, the supernatant was aspirated and replaced with 2% FBS/DMEM media containing the respective compounds at the indicated concentrations. The cells were then incubated at 37°C for 48 to 66 hours. Supernatants were collected and heat inactivated at 80°C for 20 minutes. The Luna Universal One-Step RT-qPCR Kit (New England Biolabs) was used for the detection of genomic SARS-CoV-2 RNA through RT-qPCR using a thermocycler (QuantStudio 6 thermocycler; Applied Biosystems). Specific primers targeting the N-region viral gene: Forward: 5’ TAATCAGACAAGGAACTGATTA Reverse: 5’ CGAAGGTGTGACTTCCATG were used. Cycle threshold (Ct) were determined by the second derivative maximum method in the QuantStudio software. The quantity of viral genomes is expressed as Ct and was normalized against the Ct values of the negative (PBS) and positive controls (remdesivir 25 µM). Curve fits and IC50 values were obtained in Prism using the variable Hill slope model.

### Cell viability assay

Cell viability assays were performed in compound-treated cells using the CellTiter-Glo assay according to the manufacturer’s instructions (Promega) and a Berthold Centro XS LB960 luminometer. 3,000 cells/well of Vero E6 were seeded in white with clear bottom 384-well plates. The following day, compounds were added at concentrations indicated. PBS only and 10 μM camptotecin (Sigma-Aldrich) controls were added in each plate. After 48 h incubation, 10 μl of CellTiter Glo reagent was added in each well and the luminescence was recorded using a luminometer (Berthold Technologies) with 0.5 sec integration time. Raw data were normalized against appropriate negative (0 %) and positive controls (100 %) and are expressed in % of viral replication inhibition or % of cytotoxicity.

### Immunofluorescence imaging of primary human nasal epithelium cells and K18 mouse tissues

MucilAir^TM^, reconstructed human nasal epithelial cells (hNECs), previously differentiated for 4 weeks, were cultured in 700 µL MucilAir^TM^ media on the basal side of the air/liquid interface (ALI) cultures as described^65^. For imaging, cells were fixed on the apical and basal sides with 4% PFA for 15 minutes. Cells were stained with anti-ACE2 VHH-B07-Fc at 6 µg/ml revealed with a Goat anti-Human Alexa Fluor-488 secondary Ab (Jackson Immuno Research Laboratories), Phalloidin Atto 565 (Sigma-Aldrich) (F-actin staining) or α-tubulin (ThermoFisher Scientific) revealed with a Goat anti-Rabbit Alexa Fluor 555 secondary Ab (Invitrogen), and DAPI (ThermoFisher Scientific, 1mg/ml) and imaged using the LSM-700 confocal microscope (Zeiss) as described^65^ ^33^.

For immunofluorescence experiments on whole-mount preparations of mouse organs, following euthanasia, samples of nasal turbinates and lungs of K18-hACE2 mice were collected and fixed in phosphate-buffered saline (PBS) containing 4% paraformaldehyde for 24h at 4°C. Then, nasal respiratory epithelium, olfactory epithelium and lung bronchioli were finely excised and post-fixed in PBS containing 4% paraformaldehyde for 1 hour, rinsed in PBS and immersed in blocking buffer (20% normal goat serum (NGS) and 0.3% Triton X-100 in PBS) for 1 hour at room temperature, and then incubated overnight at 4 °C in primary antibodies (in PBS containing 2% bovine serum albumin (BSA)) : VHH-B07-Fc (at 6 µg/ml) and rat anti-αtubulin (1/200; ThermoFisher Scientific). After rinsing in PBS, samples were incubated for 2 hours at room temperature in PBS-2% BSA with the appropriate secondary antibodies (goat anti-human Alexa Fluor 547 (Jackson Immuno Research Laboratories) ; donkey anti-rat Alexa-Fluor 488 (Invitrogen)), Phalloidin Atto 647 (Sigma-Aldrich) (F-actin staining) and DAPI (TermoFisher Scientific, 1mg/ml). Fluorescent immunolabellings were analyzed with a Zeiss LSM-700 confocal microscope.

### SARS-CoV-2 infection and treatment in K18-hACE2 mouse model

B6.Cg-Tg(K18-ACE2)2Prlmn/J mice (stock #034860) were imported from The Jackson Laboratory and bred at the Institut Pasteur under strict specific pathogen-free conditions. Infection studies were performed on 10 to 13 wk-old male and female mice, in animal biosafety level 3 (BSL-3) facilities at the Institut Pasteur. All animals were handled in strict accordance with good animal practice. Animal work was approved by the Animal Experimentation Ethics Committee (CETEA 89) of the Institut Pasteur (project dap 210050), and authorized by the French Ministry of Research (under project 31816) before the experiments were initiated. Twenty-four hours before infection, anesthetized (ketamine/xylazine) mice were administered intra-nasally (i.n.) with either 0.9% NaCl (40 µl), control VHH (7 mg/kg in 40 µl) or B07-Fc (7 mg/kg in 50 µl). On the next day, they were again anesthetized (ketamine/xylazine) and inoculated i.n. with 1 × 10^5^ PFU of SARS-CoV-2 XBB.1.5 (10 μl/nostril). On day 3 post-infection, mice were euthanized by ketamine/xylazine overdose. The trachea and upper lung lobes were fixed by submersion in 10% phosphate buffered formalin for 24-36 hours prior to removal from the BSL3 for processing and transfer in 70% ethanol. Lower lung lobes (and nasal turbinates in a subset of mice) were dissected and frozen at −80°C. Samples were homogenized in 400 μl of cold PBS using lysing matrix M (MP Biomedical) and a MP Biomedical FastPrep 24 Tissue Homogenizer. Viral RNA was extracted using an extraction robot IDEAL-32 (IDsolutions) and the NucleoMag Pathogen extraction kit (Macherey Nagel). Viral RNA quantification was performed by quantitative reverse transcription PCR (RT-qPCR) using the IP4 set of primers and probe (nCoV_IP4-14059 Fw GGTAACTGGTATGATTTCG and nCoV_IP4-14146 Rv CTGGTCAAGGTTAATATAGG giving a 107bp product, and nCoV_IP4-14084 Probe(+)TCATACAAACCACGCCAGG [5’]Hex [3’]BHQ-1) and the Luna Universal Probe One-Step RT-qPCR Kit (NEB). Serial dilutions of a titrated viral stock were analyzed simultaneously to express viral loads as PFU equivalents (eqPFU) per gram of tissue.

### Measurement of VHH concentration in lungs homogeneate by ELISA

sACE2 (2 µg/mL) were coated on polystyrene plate (ThermoScientific 469949) over night at 4°C. After washing with PBS/Tween 0.1%, plates were incubated with B07-Fc (from 0.07 to 10 ng/ml) or homogeneate of lower lung lobes in PBS 0.5% Gelatin, 0.1% Tween-20 (PGT solution) for 90 min at 37°C. A negative control (sACE2 alone) was prepared under the same conditions. Then, plates were incubated at 37°C with Peroxydase Affinipure rabbit anti Human IgG (Jackson ImmunoResearch 309-035-008, 1/1000) in PTG for 45 min and treated with ortho-phenylenediamine solution containing 0.1% hydrogen peroxid (H2O2) for 5 min. The reaction was stopped by adding chlorhydric acid 3M (HCl) and the absorbance of the wells was measured at 492λ in a spectrophotometer-microplate reader (Sunrise, Tecan).

### SARS-CoV-2 infection and treatment in Syrian golden hamster

The animals used were male Syrian golden hamsters (*Mesocricetus auratus*, strain RjHan:AURA), 6 week of age (weight between 70-90 grams), obtained from Janvier Laboratories. Golden hamsters were housed and manipulated in class III safety cabinets in the Institut Pasteur animal facilities accredited by the French Ministry of Agriculture. Animal work was approved by the Animal Experimentation Ethics Committee (CETEA) of the Institut Pasteur (project dap 210011) and authorized by the French legislation (project #21045) in compliance with the European Communities Council Directives (2010/63/UE, French Law 2013-118, February 6, 2013) and according to the regulations of Institut Pasteur Animal Care Committees before the experiments were initiated. Anesthetized animals received intranasal inoculation (5 mg/kg) of VHH-B07-Fc or VHH-ctl-Fc. Twenty-four hours after inoculation, hamsters were infected intranasally (i.n.) with 10^4^ PFU of SARS-CoV-2 XBB.1.16.1 as previously described ^63^.

All hamsters were followed-up daily. At day 3 post-infection, animals were euthanized with an excess of anesthetics (ketamine, xylazine and laocaine) (AVMA Guidelines 2020), and samples collected. 100 mg lung fragments were ground in Lysing Matrix X tubes (MP Biomedicals) using a grinder at a speed of 6.5 m/s for 30 seconds and centrifuged at 2,000 g for 10 minutes (4°C) to clarify the supernatant.

For the evaluation of the number of SARS-CoV-2 copies in tissues by RT-qPCR, 100 µl of each grinding solution was mixed with 500 μL of Trizol to deactivate virulence. The total RNA was then extracted using the Direct-zol RNA MiniPrep Kit (Zymo Research) and quantified. The presence of genomic SARS-CoV-2 RNA in these samples was evaluated by one-step RT-qPCR in a final volume of 5 μl per reaction in 384-well PCR plates using a thermocycler (QuantStudio 6 thermocycler; Applied Biosystems). The primers used targeted N-region viral gene: Forward: 5’ TAATCAGACAAGGAACTGATTA Reverse: 5’ CGAAGGTGTGACTTCCATG. Viral load quantification (expressed as RNA copy number/g lung) was assessed by linear regression using a standard curve of seven known quantities of RNA transcripts containing the N-region viral gene. All collected data were plotted and analyzed using GraphPad Prism software. Mann-Whitney tests were performed for statistical analysis.

### Statistical analysis

Figures were generated using GraphPad Prism 9 (GraphPad software). Statistical analysis was conducted using GraphPad Prism 9. Statistical significance between different groups was calculated using the tests indicated in each figure legend.

## Acknowledgments

We are grateful to Julian Buchrieser (Virus and Immunity unit, Institut Pasteur) and Erika Cecon (Institut Cochin, Paris, France) for the gifts of vectors expressing ACE2 and Hugo Mouquet (Laboratory of Humoral Immunology, Institut Pasteur, Paris, France) for anti-spike mAb antibody. We acknowledge the Molecular Biophysics Platform (Institut Pasteur, Paris) for soluble proteins quality control, in particular Bertrand Raynal, Sandrine Rosario, and Sébastien Brule. We acknowledge the National Reference Centre for Respiratory Viruses hosted by Institut Pasteur (Paris, France). We thank the staff of the Animal Facility (C2RA) for the breeding and care of the K18-hACE2 mice. We acknowledge the members of the Direction des Applications de la Recherche et des Relations Industrielles (DARRI, Institut Pasteur, Paris) involved in the project, particularly Corinne Sarrazin, Serge Pauillac and Michel Perez from the patent office. We thank all the members of the Dynamics of Host-Pathogen Interactions Unit (Institut Pasteur) for help with experiments and discussions, particularly Geneviève Janvier. We thank Françoise Porrot and Florence Guivel-Benhassine (Virus and Immunity unit, Institut Pasteur, Paris) for virus production.

## Funding

Institut Pasteur “Programme Federateur de Recherche” (PFR-6-SI-COV2 project), Urgence COVID-19 Fundraising Campaign of Institut Pasteur (FR, JE, OS, AB).

The French Government’s Investissement d’Avenir program, Laboratoire d’excellence “Integrative Biology of Emerging Infectious Diseases” (grant ANR-10-LABX-62-IBEID) (XM, FR, JE, OS, AB).

Programme Fédérateur de Recherche SARS-CoV-2 & COVID-19, PFR-4 Long Covid (XM).

Laboratoire d’excellence “Milieu Interieur” (JE).

Fondation pour la Recherche Médicale (FRM) (OS).

ANRS (OS).

The Vaccine Research Institute (ANR-10-LABX-77) (OS)

ANR / FRM Flash Covid PROTEO-SARS-CoV-2 (OS).

ANR Coronamito (OS).

HERA european funding (OS).

Sanofi (OS).

IDISCOVR (OS).

ANRS-MIE (BIOVAR and PRI projects of the EMERGEN research program) (XM, FA).

SB is the recipient of an MESR/Ecole Doctorale BioSPC, Université Paris Cité, fellowship; FS is the recipient of an Erasmus program. DP is supported by the Vaccine Research Institute.

The Opera system was co-funded by Institut Pasteur and the Région île de France (DIM1Health). Funding sources are not involved in the study design, data acquisition, data analysis, data interpretation or manuscript writing.

## Author contributions

Conceptualization: AB

Proteins productions: IF, ED, PL, GA

Alpaca immunization: PL, GA

*In vitro* assays: SB, IS, FS, AB, JC, JT-R, PL, GA

*In vivo* assays: LC, EG, SG

Viral strains collection: XM, FA, OS

Imaging: VM, DP

Supervision: SP, XM, FA, FR, PL, JE, OS, AB

Writing—original draft: AB wrote the manuscript with contributions from all authors.

## Data availability

The datasets generated during the current study are available from the corresponding author on reasonable request. The raw data generated in this study are provided in the Source Data file.

## Supplementary Figure Legends

**Supplementary Fig. 1: VHHs binding on cells expressing ACE2.** (**A)** B07, B09, and B10 binding on A549-ACE2 (top) and Vero E6 (bottom) cells. Cells were incubated with the VHHs (10 µg/ml), stained with an anti-myc antibody and a AF488-conjugated anti-mouse antibody, before being analyzed by flow cytometry. Parental A549 cells or VHH IgE were used as control. Data are mean ± SD of at least 3 independent experiments. (**B)** Fluorescence diagram overlays: B07, B09, B10 efficacy on different cell lines expressing exogenous (A549-ACE2) or endogenous ACE2 (Vero E6). Background (blue) corresponds to the fluorescence intensity obtained on parental cells (A549) or using a VHH control (anti-IgE VHH) (Vero E6). (**C)** B07-Fc binding on A549-ACE2 and Vero E6 cells. Cells were incubated with 0.1 µg/ml B07-Fc and a AF488-conjugated secondary antibody and analyzed as in **A**. Data are mean ± SD of at least 3 independent experiments. (**D)** Fluorescence diagram overlays performed as in B on A549-ACE2 and Vero E6 cells.

**Supplementary Fig. 2: B07-Fc staining and toxicity.** (**A)** B07-Fc detected ACE2 on ciliated cells of human nasal epithelial cells (hNEC). Representative immunofluorescence staining of ACE2 (Red: B07-Fc staining) in combination with a-tubulin (Yellow). Scale bars: left, 10 µm; right, 5 µm. (**B)** Vero E6 cells viability in the presence of B07-Fc measured by quantitating ATP using the CellTiter-Glo assay (triplicates).

**Supplementary Fig. 3: Relative cell surface expression of hACE2 polymorphic mutants.** HEK293 cells transfected with plasmids encoding myc-hACE2-WT or the indicated mutant were stained with an antibody against the myc epitope (**A, B**) or B07-Fc (**B**) and analyzed by flow cytometry. (**A**) Bars represent staining efficiency for cells expressing myc-ACE2 mutants relative to cells expressing myc-ACE2-WT after anti-myc staining. Data are means ± SD of at least two experiments. (**B**) Fluorescence diagram overlays of myc-ACE2-E35A and myc-ACE2-H34Q relative to myc-hACE2-WT after anti-myc or B07-Fc staining. Background (Grey) corresponds to the fluorescence intensity obtained using the secondary antibody alone.

**Supplementary Fig. 4: Effect of VHH B07-Fc in hamster ACE2. (A)** Schematic diagram showing the experimental design of B07-Fc prophylaxy in XBB.1.16.1 infected hamsters. Animals received intranasally (i.n.) 5 mg/kg VHH B07-Fc (B07-Fc) or 5 mg/ml VHH-Fc ctl (ctl-Fc). Twenty four hours later, they were infected with 10^4^ PFU XBB.1.16.1 intranasally (i.n.). Three days post-infection, lungs were collected for analysis. Animal behavior and weight were followed each day. **(B)** RNA load measured by RT-qPCR of SARS-CoV-2 in lung. Data are mean ± SD of 4 animals. Mann-Witney test: P value 0.0286*.

