## Supplementary figures for "Intranasal delivery of a broadly neutralizing single domain antibody targeting ACE2 protects against SARS-CoV-2 infection"

### Supplementary Figure 1

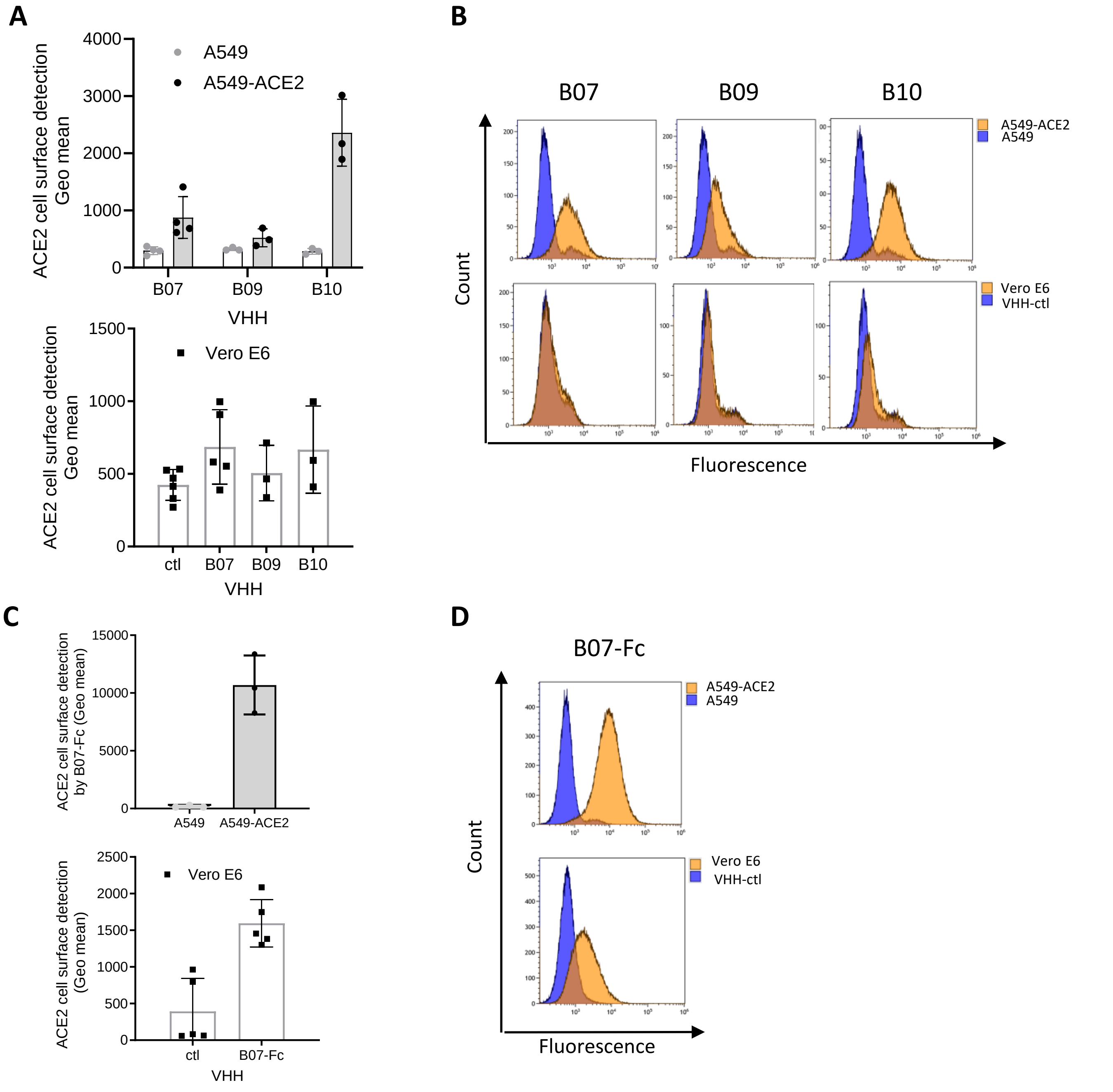


**Supplementary Fig.1: VHHs binding on cells expressing ACE2. (A)** B07, B09, and B10 binding on A549-ACE2 (top) and Vero E6 (bottom) cells. Cells were incubated with the VHHs (10 µg/ml), stained with an anti-myc antibody and a AF488-conjugated anti-mouse antibody, before being analyzed by flow cytometry. Parental A549 cells or VHH IgE were used as control. Data are mean ± SD of at least 3 independent experiments. (**B**) Fluorescence diagram overlays: B07, B09, B10 efficacy on different cell lines expressing exogenous (A549-ACE2) or endogenous ACE2 (Vero E6). Background (blue) corresponds to the fluorescence intensity obtained on parental cells (A549) or using a VHH control (anti-IgE VHH) (Vero E6). (**C**) B07-Fc binding on A549-ACE2 and Vero E6 cells. Cells were incubated with 0.1 µg/ml B07-Fc and a AF488-conjugated secondary antibody, and analyzed as in **A**. Data are mean ± SD of at least 3 independent experiments. (**D**) Fluorescence diagram overlays performed as in **B** on A549-ACE2 and Vero E6 cells.

### Supplementary Figure 2

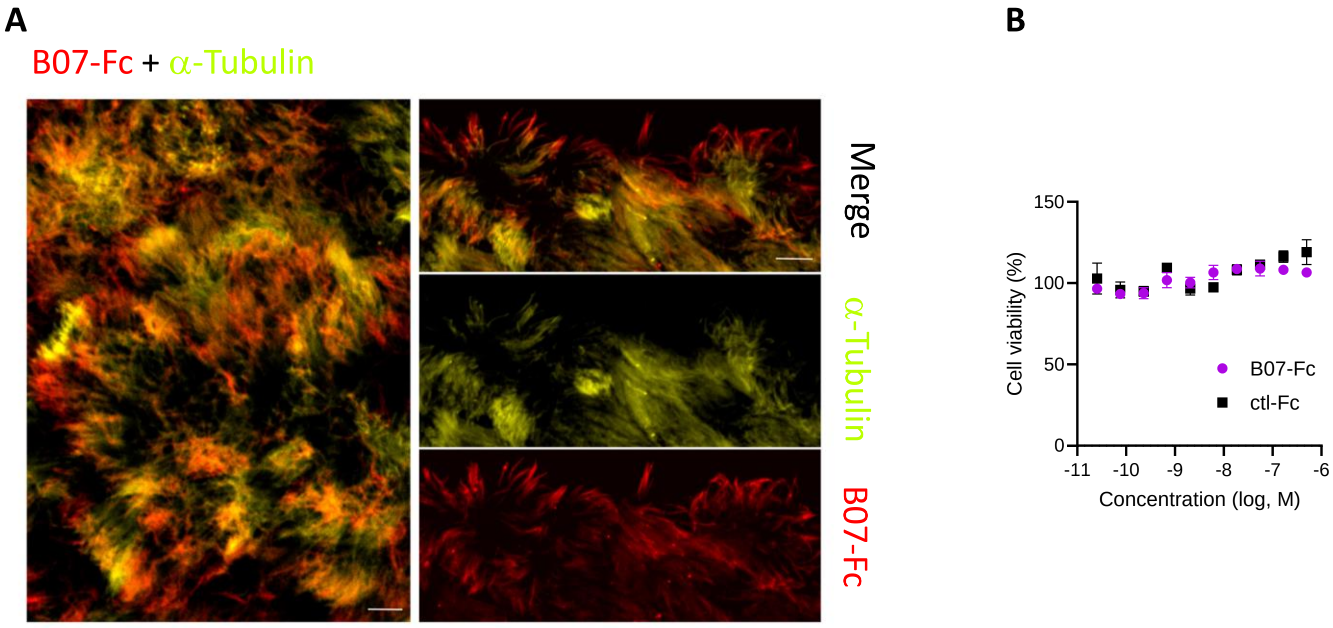


**Supplementary Fig. 2**: **B07-Fc staining and toxicity**. (**A**) B07-Fc detected ACE2 on ciliated cells of human nasal epithelial cells (hNEC). Representative immunofluorescence staining of ACE2 (Red: B07-Fc staining) in combination with $\alpha$-tubulin (Yellow). Scale bars: left, 10 µm; right, 5 µm. (**B**) Vero E6 cells viability in the presence of B07-Fc measured by quantitating ATP using the CellTiter-Glo assay (triplicates).

### Supplementary Figure 3

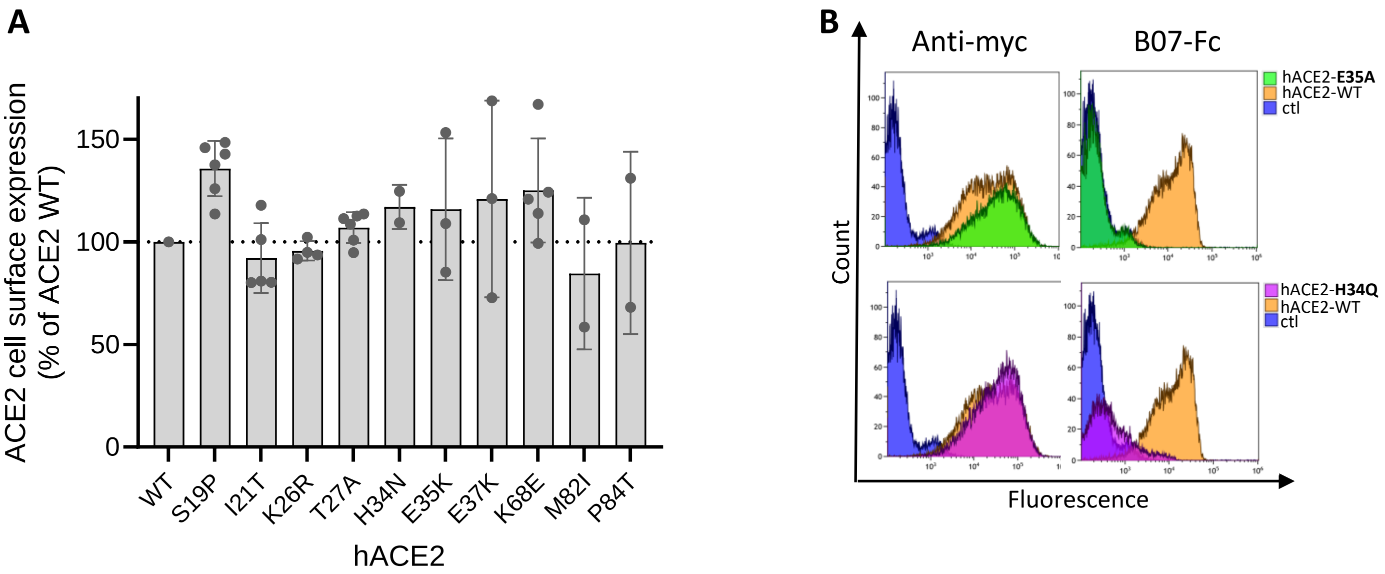


**Supplementary Fig. 3: Relative cell surface expression of ACE2 mutants.** HEK293 cells transfected with plasmids encoding myc-hACE2-WT or the indicated mutant were stained with an antibody against the myc epitope (**A, B**) or with B07-Fc (**B**) and analyzed by flow cytometry. (**A**) Bars represent staining efficiency for cells expressing myc-hACE2 mutants relative to cells expressing myc-hACE2-WT after anti-myc staining. Data are means ± SD of at least two experiments. (**B**) Fluorescence diagram overlays of myc-hACE2-E35A and myc-hACE2-H34Q relative to myc-hACE2-WT after anti-myc or B07-Fc staining. Background (Blue) corresponds to the fluorescence intensity obtained using the secondary antibody alone.

### Supplementary Figure 4

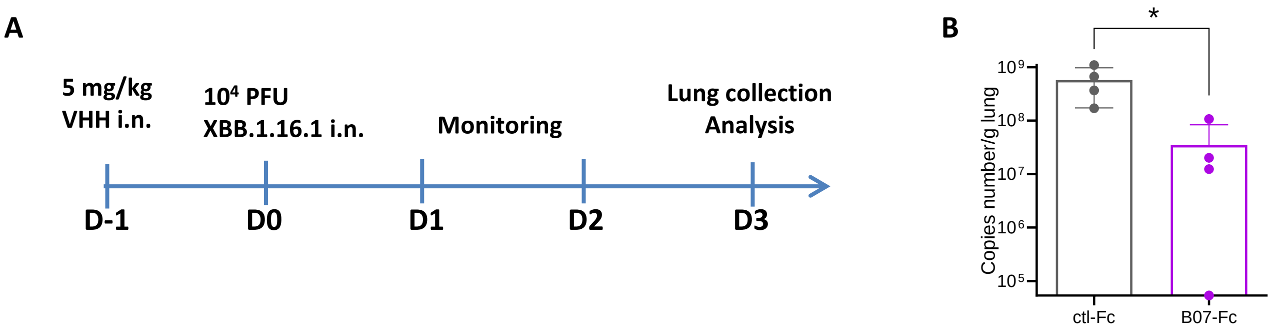


**Supplementary Fig. 4: Effect of VHH B07-Fc on hamster ACE2. (A)** Schematic diagram showing the experimental design of B07-Fc prophylaxy in XBB.1.16.1 infected hamsters. Animals received intranasally (i.n.) 5 mg/kg VHH-B07-Fc (B07-Fc) or 5 mg/ml VHH-Fc ctl (ctl-Fc). Twenty four hours later, they were infected with $10^4$ PFU XBB.1.16.1 intranasally (i.n.). Three days post-infection, lungs were collected for analysis. Animal behavior and weight were followed each day. **(B)** RNA load measured by RT-qPCR of SARS-CoV-2 in lung. Data are mean ± SD of 4 animals. Mann-Witney test: P value 0.0286*.
